# Persistent one-way walking in a circular arena in *Drosophila melanogaster* Canton-S strain

**DOI:** 10.1101/145888

**Authors:** Chengfeng Xiao, Shuang Qiu, R Meldrum Robertson

**Author notes:** Corresponding author: Chengfeng Xiao, Department of Biology, Queen’s University, Kingston, Ontario K7L 3N6, Canada, 1-613-533-6000 ext 75510, 1-613-533-6617 (fax).

## Abstract

We describe persistent one-way walking of *Drosophila melanogaster* in a circular arena. Wild-type Canton-S adult flies walked in one direction, counter-clockwise or clockwise, for minutes, whereas white-eyed mutant *w*^1118^ changed directions frequently. Locomotion in the circular arena could be classified into four components: counter-clockwise walking, clockwise walking, nondirectional walking and pausing. Genetic analysis revealed that while wild-type genetic background was associated with reduced directional change and reduced numbers of one-way (including counterclockwise and clockwise) and nondirectional walks, the *white* (*w*^+^) locus promoted persistent oneway walking by increasing the maximal duration of one-way episodes. The promoting effect of *w*^+^ was further supported by the observations that (1) *w*^+^ duplicated to the Y chromosome, (2) four genomic copies of *mini-white* inserted on the autosomes, and (3) pan-neuronal overexpression of the White protein increased the maximal duration of one-way episodes, and that RNAi knockdown of *w*^+^ in the neurons decreased the maximal duration of one-way episodes. These results suggested a pleiotropic function of *w*^+^ in promoting persistent one-way walking in the circular arena.

## Introduction

Walking locomotion in *Drosophila melanogaster* (fruitfly) displays many distinctive features. Negative geotaxis and positive phototaxis are the classical stereotypes of locomotion in adult flies (Carpenter 1905; McEwen 1918; Cole 1922). Directional persistence and local wall attraction are the two main features of walking in open field arenas (Soibam et al. 2012). Wing-clipped flies walk back and forth towards two dark strips that are inaccessible, oppositely positioned and relatively narrow in Buridan’s paradigm (Colomb et al. 2012). In addition, adult flies are unwilling to walk through confined spaces, a phenomenon termed claustrophobia (Ewing 1963). While restricted in small arenas, flies perform relentless walking for hours (Cole 1995; Xiao and Robertson 2015).

A common interest is to understand the genetic basis for walking behavior in *Drosophila*. The wild-type Canton-S and white-eyed mutant *w*^1118^ flies have different walking performance observed in several experimental settings. Canton-S flies walk towards light more often than *w*^1118^ flies (Kain et al. 2012). In circular arenas, Canton-S flies have higher boundary preference than *w*^1118^ flies (Liu et al. 2007; Xiao and Robertson 2015; Qiu et al. 2017). Canton-S recover walking after anoxia faster and more consistently than *w*^1118^ flies (Xiao and Robertson 2016). When rigidly restrained, Canton-S flies show spontaneous and rhythmic motor activities that can be recorded extracellularly from the brain, whereas *w*^1118^ flies have greatly reduced rhythmic motor activities (Qiu et al. 2016). These findings raise a concern: either the *white* (*w*^+^) gene, which is null-mutated in *w*^1118^ flies, or its genetic background, is responsible for the walking performance of Canton-S different from *w*^1118^ flies.

*w*^+^ is a classic eye-color gene discovered by Thomas Hunt Morgan in 1910 (Morgan 1910). The product of *w*^+^ is a subunit of transmembrane ATP-binding cassette (ABC) transporter, which loads vesicles*/*granules with biogenic amines (Borycz et al. 2008), second messenger (Evans et al. 2008), metabolic intermediates (Sullivan and Sullivan 1975; Anaka et al. 2008) and pigment precursors (Sullivan and Sullivan 1975; O’Hare et al. 1984; Dreesen et al. 1988; Tearle et al. 1989). Increasing evidence has supported the proposal that *w*^+^ possesses pleiotropic housekeeping functions in addition to eye pigmentation (Zhang and Odenwald 1995; Hing and Carlson 1996; Campbell and Nash 2001; Evans et al. 2008; Borycz et al. 2008; Anaka et al. 2008; Xiao and Robertson 2016, 2017; Xiao et al. 2017). We hypothesized that *w*^+^ modulates locomotor behavior and promotes persistent walking performance.

In this study, we describe persistent one-way walking of flies in the circular arenas. Wild-type Canton-S flies walk in one direction, counter-clockwise or clockwise, for minutes, whereas *w*^1118^ flies change directions frequently. We extract the behavioral components of walking in the arena, and show that counter-clockwise walking and clockwise walking are the two main locomotor components, and that sporadic pausing is less associated with directional change than with directional persistence. We further show that while wild-type genetic background reduces the number of directional changes, *w*^+^ promotes persistent one-way walking by increasing the maximal duration of one-way walking.

## Materials and methods

### Flies

Flies used in this study and their sources were: Canton-S (Bloomington *Drosophila* Stock Center (BDSC) # 1); *w*^1118^ (L. Seroude laboratory); Oregon-R (BDSC # 2376); Hikone-AS (BDSC # 3); Florida-9 (BDSC # 2374); *w*^1^ (BDSC # 145); *w*^*a*^ (BDSC # 148); *w*^*c*^ ^*f*^ (BDSC # 4450); UAS-bPDE5 (Vermehren-Schmaedick et al. 2010); UAS-*white* (D4) and UAS-*white* (H8) (Evans et al. 2008); UAS-Httex1-Qn-eGFP (n = 23, 72 or 103) (Zhang et al. 2010); 10*×*UAS-IVS-mCD8::GFP (attP40) (BDSC # 32186); 10*×*UAS-IVS-mCD8::GFP (attP2) (BDSC # 32185); 10*×*UAS-IVSGFP-WPRE (attP2) (BDSC # 32202); UAS-mito-HA-GFP.AP (BDSC # 8442); UAS-mito-HAGFP.AP (BDSC # 8443); tubP-Gal80^*ts*^ (BDSC # 7019); NP6520-Gal4 (Awasaki et al. 2008); elavGal4 (BDSC # 8765) and UAS-*white*-RNAi (BDSC # 31088). *w*^+^ F10 and *w*^1118^ F10 were the fly lines generated by a serial backcrossing between Canton-S and *w*^1118^ flies for ten generations (Xiao and Robertson 2015). *w*^+^ duplication lines (*w*^1118^/Dp(1;Y)*B*^*S*^*w*^+^*y*^+^ and *w*^1118^/Dp(1;Y)*w*^+^*y*^+^) were generated previously (Xiao and Robertson 2016; Xiao et al. 2017). Flies were maintained with standard medium (cornmeal, agar, molasses and yeast) at 21-23°C with 60-70 % relative humidity. An illumination of light*/*dark (12*/*12 h) cycle was provided with three light bulbs (Philips 13 W compact fluorescent energy saver) in a room around 133 square feet. Flies were collected within 0 - 2 days after emergence. We used pure nitrogen gas to anesthetize flies during collection time. Collected flies were raised in food vials at a density of 20 - 25 flies per vial for at least three additional days. A minimum of three days free of nitrogen exposure was guaranteed before the test. The ages of tested flies were 4 - 9 days old. Unless otherwise indicated, male flies were used for experiments. To avoid natural peak activities in the mornings and evenings (Grima et al. 2004), experiments were performed during the light time with three hours away from light on*/*off switch.

### Locomotor assay

Locomotor assay was performed by following a reported protocol (Xiao and Robertson 2015). In general, flies were loaded into circular arenas (1.27 cm diameter and 0.3 cm depth) with one fly per arena. The depth of 0.3 cm was considered to allow flies to turn around but suppress vertical movement. We machined 128 arenas (8 *×* 16) in an area of 31.0 cm *×* 16.0 cm Plexiglas. The bottom side of arena was covered with thick filter paper allowing air circulation. The top was covered by a slidable Plexiglas with holes (0.3 cm diameter) at one end for fly loading. The Plexiglas with arenas was secured in a large chamber (48.0 cm *×* 41.5 cm *×* 0.6 cm). A flow of room air (2 L/min) was provided to remove the effect of dead space (Bouhuys 1964). Illumination for the settings was provided by using a white light box (Logan portaview slide*/*transparency viewer) with a 5000 K (“daylight”) color-corrected fluorescent lamp. Dim red illumination was generated by covering a red filter (Roscolux #26 Light Red, Rosco Canada) on the light box. A time of 5-min was allowed for flies to adapt to the experimental settings. Locomotor activities were video-captured at 15 frames per second, and stored for post analysis. Fly positions (the locations of center of mass) with 0.2 s interval were computed by custom-written fly tracking software (Xiao and Robertson 2015). For each fly, a dataset containing 1500 positional coordinates, from 300 s locomotion, was collected. These data were used for subsequent behavioral analysis, including the construction of time-series of 3D trajectory and the extraction of behavioral components of walking.

### Construction of 3D walking trajectory

Time-series of 3D walking trajectory was constructed by using the X-Y positions over a period of time. The function Cloud() from an R package “Lattice” (Sarkar 2008) was used for 3D data visualization. The arena size was remained unchanged throughout this study. Therefore, for simplification, we omitted the *x*, *y* axes (representing position coordinates) and *z* axis (representing time), and provided a color key as an indicator of time. For clarity, some images present only part of the data (e.g. data from 60 s locomotion).

### Extraction of behavioral components of walking

We used two parameters, walking direction (counter-clockwise or clockwise) and step size, to categorize the behavioral components of walking. Walking direction was computed using the center of arena as a reference. Computation of walking direction was performed by following these three steps: (1) calculate angular coordinates of fly positions by a trigonometric function atan2(y, x). (2) compute parameter *ω* - the angular displacement per 0.2 s. To avoid the big jump of *ω* value due to discontinuous radian rotation, we calculated *ω* twice using radian interval (0, 2π] and (-*π*, *π*], and chose one with smaller absolute value. (3) determine the walking direction as counter-clockwise (*ω >* 0) or clockwise (*ω <* 0).

We defined “counter-clockwise walking” as at least five consecutive steps with *ω >* 0, and “clockwise walking” as at least five consecutive steps with *ω <* 0. Categorized data were further rounded by allowing 1-2 steps of pausing or backward walking (with radian *<* 0.21, equivalent to 12°) without an apparent change of direction.

To improve the estimation, we separated pausing steps from active walking. Briefly, the step size (euclidean distance per 0.2 s) was computed. “pausing” was defined as at least five consecutive steps with step size *<* 0.28 mm, a distance equivalent to that between two adjacent diagonal pixels. The activities that were not categorized to “counter-clockwise walking", “clockwise walking” or “pausing” were assigned as “nondirectional walking", representing non-pausing activities with no consistent directionality, namely, activities failed to meet the criterion of a minimum of five consecutive steps in either direction.

### Analysis of walking directions before and after a pause

Steps for the analysis of walking directions before and after a pause were: (1) identify the behavioral component for the last position before a pause; (2) identify the behavioral component for the first position after a pause; (3) compare the difference between. If these two components were the same, it was deemed that this pausing was associated with directional persistence. Otherwise, the pausing was associated with directional change.

### Quantification of directional change

A directional change is defined as a switch between any two of the three walking components (i.e. counter-clockwise walking, clockwise walking and nondirectional walking). The number of directional changes was quantified as the total number of three walking components minus one. For the simplification of behavioral analysis, we omitted pausing because of its weak association with directional change.

### Statistics

Data processing and visualization were conducted by using software R (R Core Team 2014) and these supplementary packages: gdata, lattice (Sarkar 2008) and adehabitatLT (Calenge 2006). Data normality was examined by D’Agostino & Pearson omnibus normality test. Nonparametric tests (Mann-Whitney test, Wilcoxon matched pairs test and Friedman test with Dunn’s multiple comparison) were performed for the comparison of medians between groups. Data are presented as scattered dots and median. A *P <* 0.05 was considered statistically significant.

## Results

### Persistent one-way walking of wild-type flies in circular arenas

A male, wild-type Canton-S fly walked in one direction, counter-clockwise or clockwise, for minutes in a circular arena (1.27 cm diameter 0.3 cm depth). This persistent one-way walking was consistent between individuals (video S1). Male flies of a white-eyed mutant *w*^1118^, however, changed directions frequently and failed to maintain the walking direction for at least a minute (video S2). 3D walking trajectories of Canton-S flies, represented as a time-series of connected X-Y positions per 0.2 s, showed a regular, coil-like shape during locomotion (Figure 1a). The trajectories of *w*^1118^ flies displayed a shape of collapsed coil with visually increased irregularity (Figure 1b). The 2D path of Canton-S showed a strong preference for the perimeter of arena, whereas during the same period the 2D path of *w*^1118^ flies displayed reduced preference for perimeter and frequent crossing of the central region. Persistent one-way walking was also observed in Canton-S females compared with *w*^1118^ females (Figure S1). In addition, persistent one-way walking in Canton-S flies was unchanged by dim red illumination (Figure S2), which influences fly vision (McEwen 1918). Several additional wild-types, including Oregon-R, Hikone-AS and Florida-9, showed a similar performance of persistent one-way walking in circular arenas to Canton-S. Several white-eyed mutants (*w*^1^, *w*^*a*^ and *w*^*c*^ ^*f*^) displayed irregular trajectories similar to *w*^1118^ flies (Figure S3). Therefore, wild-type flies showed persistent one-way walking in the circular arenas.

**Figure 1:**
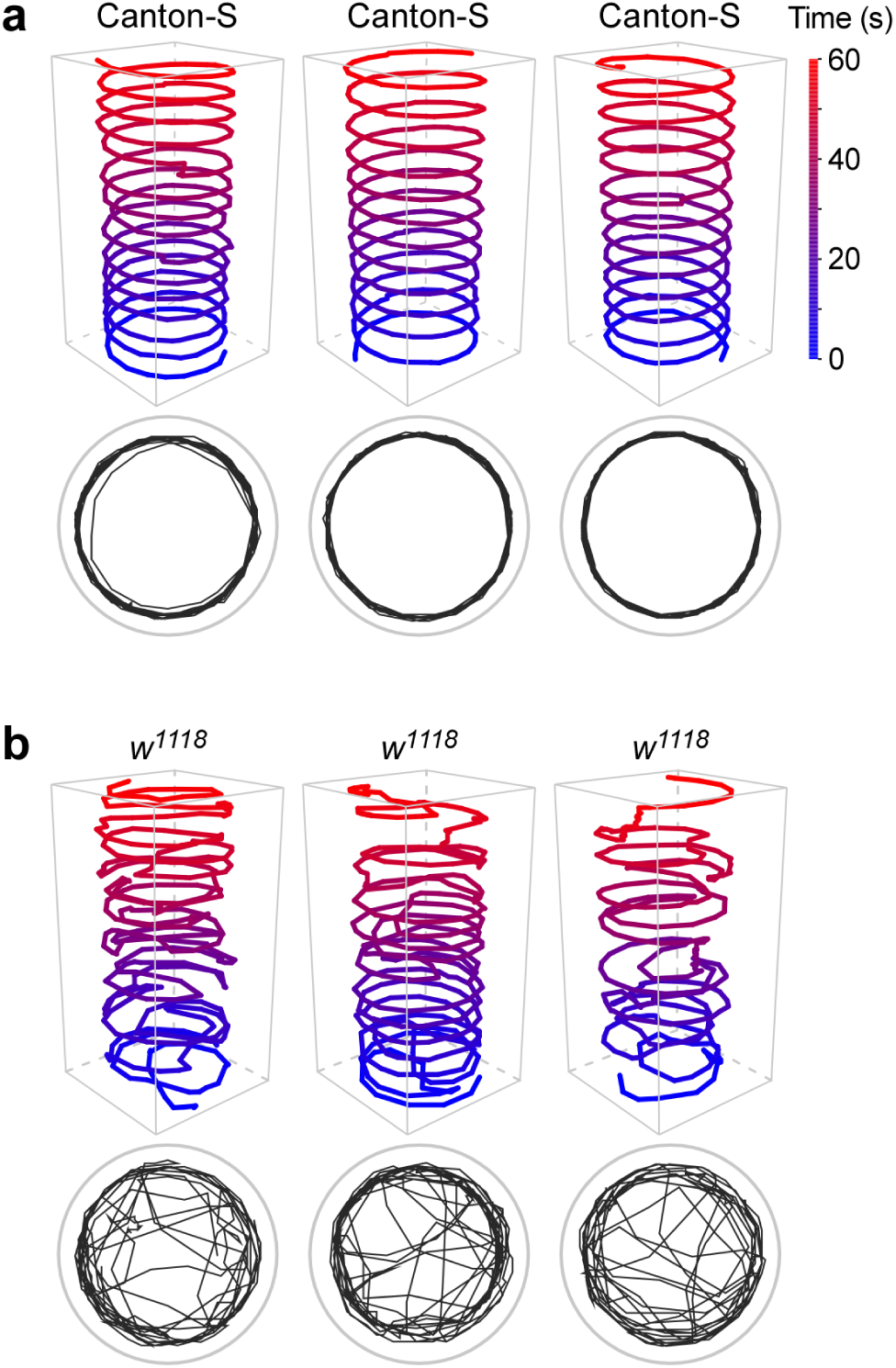
Persistent one-way walking of Canton-S flies in the circular arenas. **(a)** 3D walking trajectories and 2D path of Canton-S flies in the circular arenas (1.27 cm diameter 0.3 cm depth). There is only one fly in each arena. Color key indicates the time. **(b)** 3D walking trajectories and 2D path of *w*^1118^ flies. For clarity, walking activities from only 60 s locomotion are shown.

### Counter-clockwise walking and clockwise walking were the main components of locomotion in a circular arena

Using a fly-tracking protocol (Xiao and Robertson 2015) and software R (R Core Team 2014), we computed the parameters of locomotion and extracted the components of walking behavior.

Four walking components could be identified in a fly. They were counter-clockwise walking, clockwise walking, nondirectional walking and pausing. Canton-S showed a walking pattern that was clearly recognizable. Several flies walked in one direction, interspaced with a few pauses, throughout 300 s. In contrast, *w*^1118^ fly displayed a complicated pattern with frequent switches between walking components (Figure 2a).

**Figure 2:**
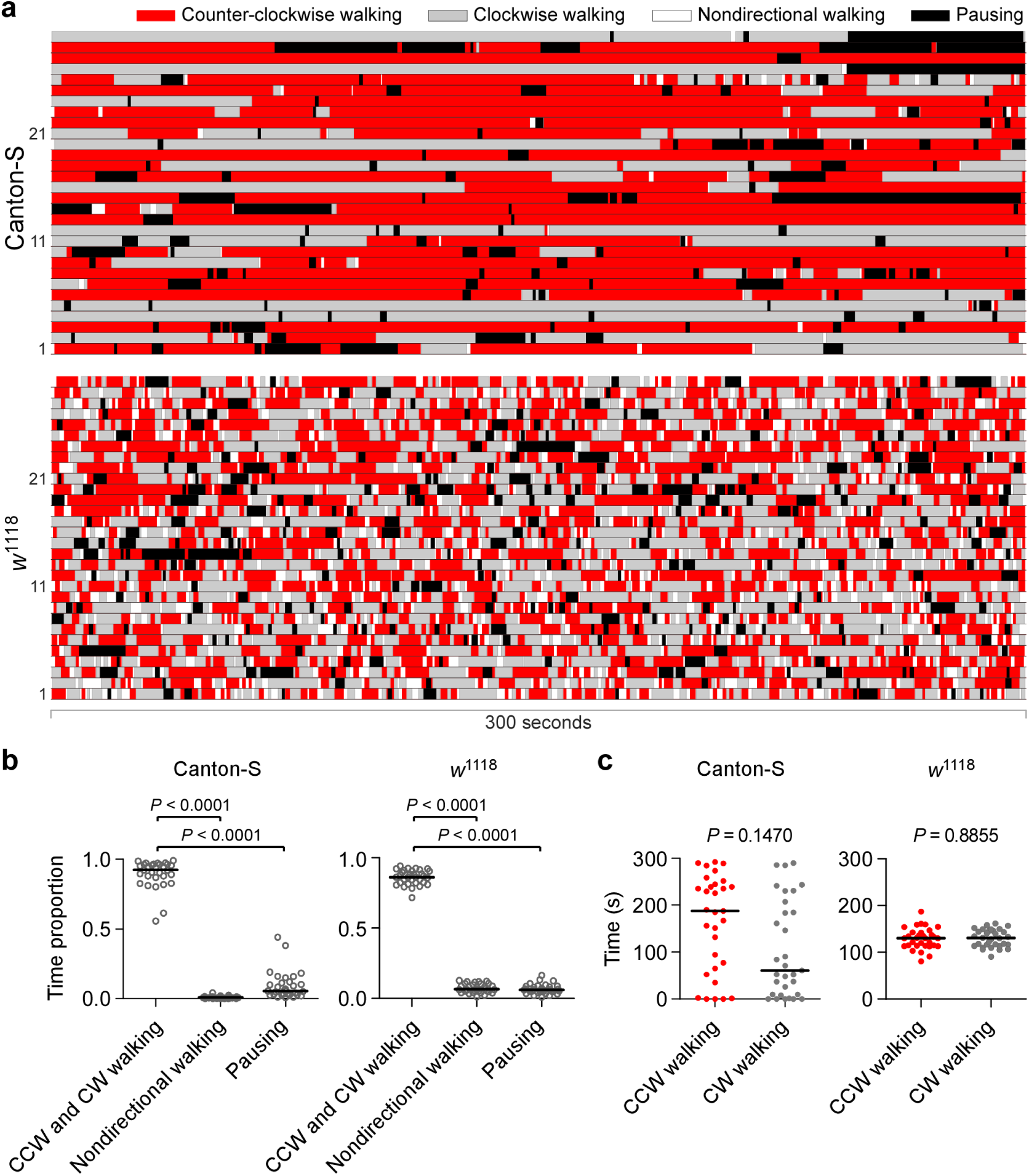
Walking components of flies in the circular arenas. **(a)** Schematic components of walking during 300 s locomotion. Shown are the counter-clockwise walking (red), clockwise walking (grey), nondirectional walking (white) and pausing (black) from 30 Canton-S and 30 *w*^1118^ flies. **(b)** Time proportions for walking components in Canton-S and *w*^1118^ flies. Data are presented as scattered dots and median (black line). CCW, counter-clockwise; CW, clockwise; *P* values from Friedman test with Dunn’s multiple comparison. **(c)** Times for CCW walking (red) and CW walking (grey) during 300 s locomotion in Canton-S and *w*^1118^ flies. *P* values from Wilcoxon matched pairs test.

Counter-clockwise walking and clockwise walking were the two main components, comprising the most time proportion in Canton-S (median 0.93, interquartile range (IQR) 0.87 - 0.97, n = 31) (Friedman test with Dunn’s multiple comparison) as well as *w*^1118^ flies (median 0.86, IQR 0.82 - 0.91, n = 32) (Friedman test with Dunn’s multiple comparison) compared respectively with other walking components. Nondirectional walking comprised a small proportion of time in Canton-S (median 0.009, IQR 0.005 - 0.013, n = 31) and *w*^1118^ flies (median 0.07, IQR 0.05 - 0.10, n = 32). The nondirectional walking included activity that failed to meet a criterion: a minimum of five consecutive steps in counter-clockwise or clockwise direction. Time proportion for pausing in Canton-S (median 0.06, IQR 0.03 - 0.12, n = 31) and *w*^1118^ flies (median 0.06, IQR 0.05 - 0.08, n = 32) was also small (Figure 2b). Compared with *w*^1118^ flies, Canton-S had increased time proportion for counter-clockwise and clockwise walking (*P* = 0.0020, Mann-Whitney test), decreased time proportion for nondirectional walking (*P <* 0.0001, Mann-Whitney test) and the same time proportion for pausing (*P* = 0.9397, Mann-Whitney test). During 300 s locomotion, the time for counter-clockwise walking (median 187.6 s, IQR 65.4 - 244.8 s, n = 31) and the time for clockwise walking (median 60.6 s, IQR 9.0 - 207.4 s, n = 31) were statistically the same in Canton-S flies (*P* = 0.1470, Wilcoxon matched pairs test). The time for counter-clockwise walking (median 130.4 s, IQR 114.3 - 142.7 s, n = 32) and the time for clockwise walking (median 130.7 s, IQR 114.4 - 146.3 s, n = 32) were also the same in *w*^1118^ flies (*P* = 0.8855, Wilcoxon matched pairs test) (Figure 2c). There was no preference for counter-clockwise or clockwise direction in circular arenas in either Canton-S or *w*^1118^ flies.

Based on this classification, the observation of persistent one-way walking in Canton-S flies could be interpreted as below: “one-way walking” denotes primarily counter-clockwise walking and clockwise walking, while “persistent” indicates the duration of a single episode of walking.

### Pausing was less associated with directional change

It was observed that flies paused sporadically in the circular arenas (see Figure 2a). It is possible that a fly pauses and changes walking direction. We examined whether the pausing was associated with directional change or directional persistence of flies in the arenas.

Canton-S flies paused several times during a period of 100 s (duration of analysis reduced for clarity). There was no obvious change of walking direction before and after a pause. Similarly, *w*^1118^ flies showed no apparent difference of walking direction before and after a pause (Figure 3a). Within 300 s, the number of pauses with directional change (median 0, IQR 0 - 1, n = 31) was remarkably lower than that with directional persistence (median 4, IQR 3 - 6, n = 31) in Canton-S (*P <* 0.0001, Wilcoxon matched pairs test) (Figure 3b). The number of pauses with directional change (median 0, IQR 0 - 2, n = 32) was also lower than that with directional persistence (median 7, IQR 4 - 8, n = 32) in *w*^1118^ flies (*P <* 0.0001, Wilcoxon matched pairs test) (Figure 3c). Thus, pausing was less associated with directional change than with directional persistence during walking in the arena.

**Figure 3:**
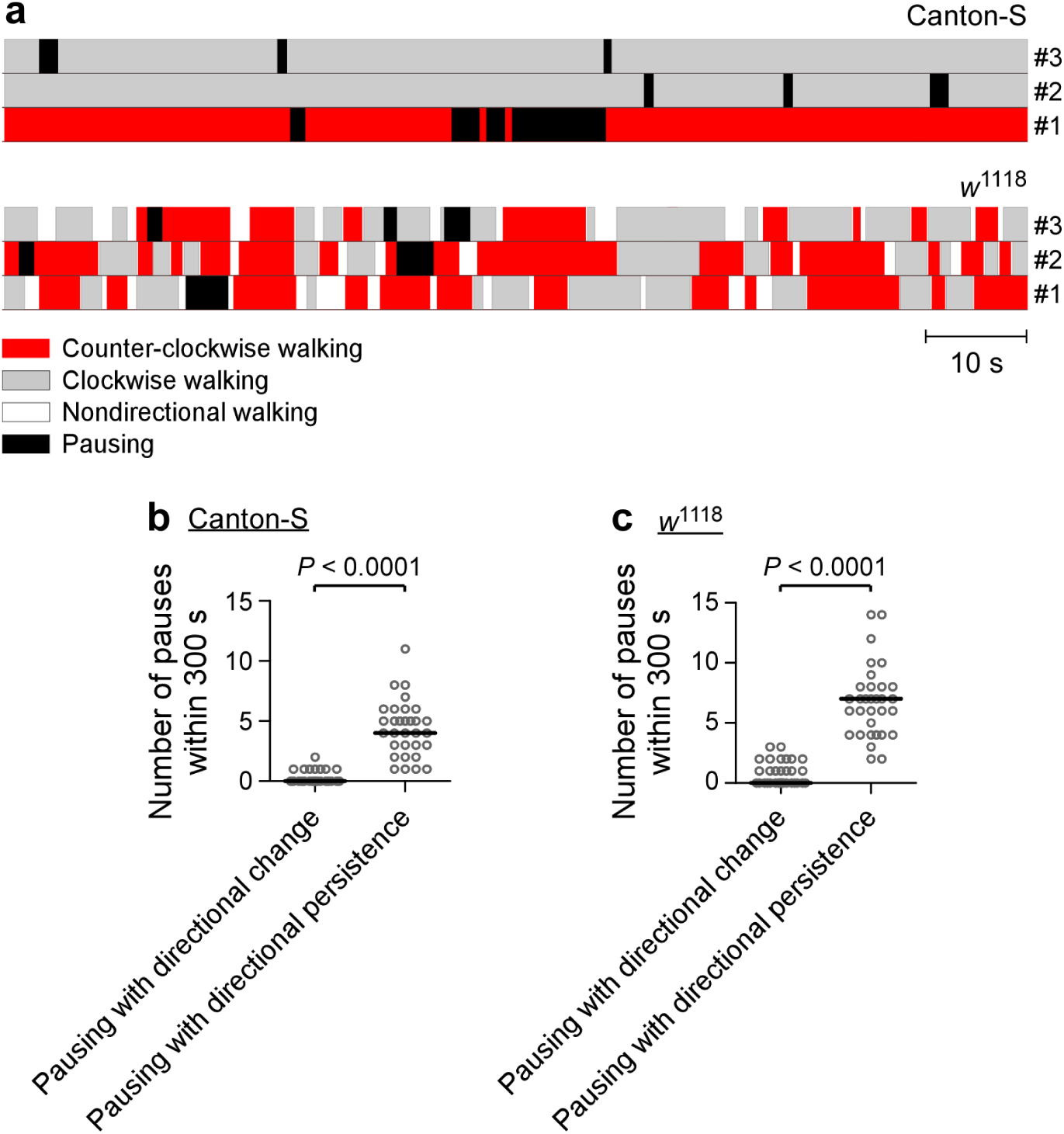
Pausing was less associated with directional change. **(a)** Schematic association between pausing and other walking components. Walking components are listed. For clarity, activities from only 100 s are shown. A time bar (10 s) is provided. **(b)** Association between pausing and directional change*/*persistence in Canton-S flies. *P* value from Wilcoxon matched pairs test. **(c)** Association between pausing and directional change/persistence in *w*^1118^ flies. *P* value from Wilcoxon matched pairs test.

### Wild-type genetic background reduced directional change of walking

For the simplification of behavioral analysis, we focused on the walking components (i.e. counterclockwise, clockwise and nondirectional walking) and explored their genetic basis. More specifically, we quantified the number of directional changes of walking in the arena. Pausing was excluded because of its weak association with directional change.

Canton-S and *w*^1118^ flies have different alleles of *w* and genetic background. We examined the contributions of *w*^+^ and its genetic background to directional change. We first compared the numbers of directional changes in *w*^+^ F1 and *w*^1118^ F1 male flies. *w*^+^ F1 was the progeny of *w*^1118^ males and Canton-S females, and reciprocally, *w*^1118^ F1 the progeny of Canton-S males and *w*^1118^ females. These two types of males had different X chromosomes (including *w* allele) and the same combinations on the second, third and fourth chromosome pairs, thus roughly the same genetic background. Both *w*^+^ F1 and *w*^1118^ F1 flies changed directions frequently in circular arenas during 300 s locomotion (Figure 4a). The number of directional changes in *w*^+^ F1 (median 51.5, IQR 44.5 - 68.5, n = 24) was comparable with that in *w*^1118^ F1 (median 57.5, IQR 39.3 - 65.5, n = 32) (*P* = 0.9011, Mann-Whitney test) (Figure 4b). Therefore, flies with roughly the same genetic background had the same number of directional changes, indicating that the genetic background but not the *w* allele was responsible for directional change of walking in the arena.

**Figure 4:**
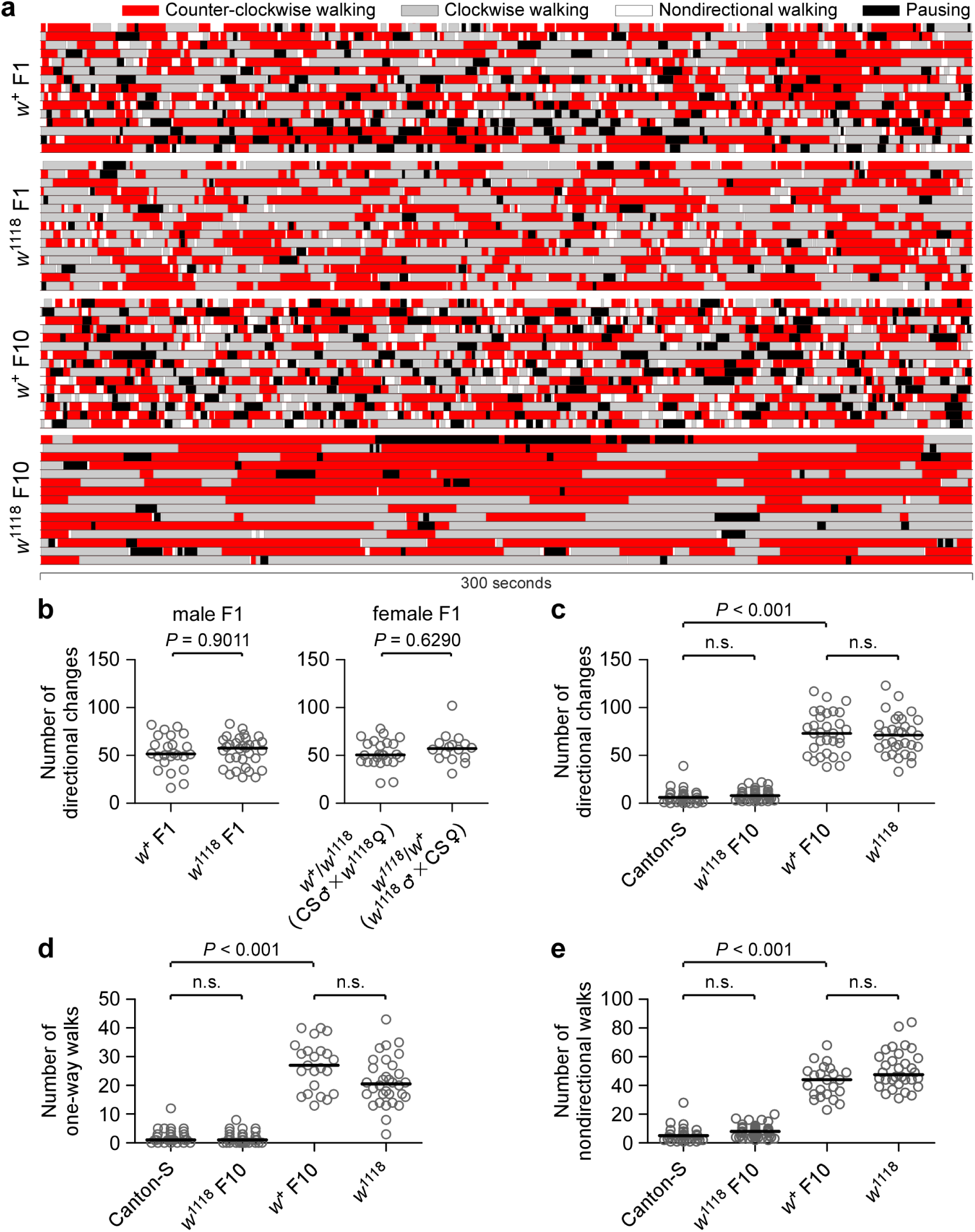
Wild-type genetic background reduces directional change and numbers of one-way and nondirectional walks. (**a**) Schematic walking components during 300 s locomotion of flies in circular arenas. Walking components and fly genotypes are indicated. Shown are the representations of 15 flies of each genotype. (**b**) Number of directional changes in male and female F1 flies. Female F1 genotypes and their parental crosses are indicated. *P* values from Mann-Whitney test. (**c**) Number of directional changes in Canton-S, *w*^1118^ F10, *w*^+^ F10 and *w*^1118^ flies. (**d**) Number of one-way walks in four groups of flies. (**e**) Number of nondirectional walks in four groups of flies. Data are presented as scattered dots and median (black line). *P* values (for **c, d and e**) from Kruska-Wallis test with Dunn’s multiple comparison; n.s. non-significance.

Female F1 flies were also examined. The number of directional changes in *w*^+^/*w*^1118^ flies (progeny of Canton-S males and *w*^1118^ females, median 50.5, IQR 43.3 - 62.8, n = 24) was statistically the same as that in *w*^1118^/*w*^+^ (progeny of *w*^1118^ males and Canton-S females, median 57.0, IQR 47.3 - 62.8, n = 16) (*P* = 0.6290, Mann-Whitney test) (Figure 4b). Thus, female F1 also changed directions frequently in circular arenas. These two types of females carried the same genomic contents (heterozygous *w* alleles and 1:1 mixed genetic background). Data supported that genetic background was associated with the number of directional changes. Additionally, these results suggested that the effect of cytoplasmic background (or maternal effect) was negligible.

We further examined the numbers of directional changes in Canton-S, *w*^1118^ F10, *w*^+^ F10 and *w*^1118^ male flies. *w*^1118^ F10 and *w*^+^ F10 were generated by a serial backcrossing between CantonS and *w*^1118^ for ten generations (Xiao and Robertson 2016). *w*^1118^ F10 flies, which carried *w*^1118^ allele in the wild-type genetic background, had the number of directional changes (median 8.0, IQR 5.0 - 11.0, n = 43) the same as Canton-S (median 6.0, IQR 1.0 - 9.0, n = 31) (*P >* 0.05, Kruskal-Wallis test with Dunn’s multiple comparison). *w*^+^ F10 flies, which carried *w*^+^ in the isogenic background, however, had the number of directional changes (median 73.0, IQR 52.0 - 94.0, n = 31) markedly greater than Canton-S (*P <* 0.001, Kruskal-Wallis test with Dunn’s multiple comparison) (Figure 4c). Therefore, a replacement of wild-type genetic background with isogenic beckground increased the number of directional changes, whereas a replacement of *w*^+^ locus with *w*^1118^ locus had no effect. Additionally, *w*^+^ F10 had the same number of directional changes as *w*^1118^ flies (median 71.0, IQR 58.3 - 85.8, n = 32) (*P >* 0.05, Kruskal-Wallis test with Dunn’s multiple comparison). Both *w*^+^ F10 and *w*^1118^ flies carried the isogenic background. These data confirmed that genetic background was responsible for the number of directional changes. The wild-type genetic background potently reduced directional change of walking in the arena.

### Wild-type genetic background reduced numbers of one-way and nondirectional walks

The directional change was reflective of transition frequency among three walking components, and was thus unable to represent the specific characterizations of individual components. We next examined the contributions of *w*^+^ and its genetic background to the numbers of one-way and nondirectional walks in the arena. Counter-clockwise walks and clockwise walks were pooled and analyzed together as one-way walks, because flies had no preference for counter-clockwise or clockwise direction.

During 300 s locomotion, the numbers of one-way walks in *w*^1118^ F10 flies (median 1, IQR 0 - 2, n = 43) were the same as Canton-S flies (median 1, IQR 1 - 3, n = 31) (*P >* 0.05, Kruskal-Wallis test with Dunn’s multiple test). The numbers of one-way walks in *w*^+^ F10 flies (median 27, IQR 17 - 32, n = 23) were markedly higher than Canton-S flies (*P <* 0.001, Kruskal-Wallis test with Dunn’s multiple test). There was no statistical difference of the number of one-way walks between *w*^+^ F10 flies and *w*^1118^ flies (median 20.5, IQR 16.3 - 26.8, n = 32) (*P >* 0.05, Kruskal-Wallis test with Dunn’s multiple test) (Figure 4d). Data indicated that the wild-type genetic background but not the *w*^+^ locus reduced the number of one-way walks.

Within 300 s locomotion, the numbers of nondirectional walks in *w*^1118^ F10 flies (median 8, IQR 4 - 10, n = 43) were the same as Canton-S flies (median 5, IQR 2 - 8, n = 31) (*P >* 0.05, Kruskal-Wallis test with Dunn’s multiple test). The numbers of nondirectional walks in *w*^+^ F10 flies (median 44, IQR 34 - 50, n = 23) were remarkably higher than Canton-S flies (*P <* 0.001, Kruskal-Wallis test with Dunn’s multiple test). There was no statistical difference of the number of nondirectional walks between *w*^+^ F10 flies and *w*^1118^ flies (median 47.5, IQR 41.8 - 59.8, n = 32) (*P >* 0.05, Kruskal-Wallis test with Dunn’s multiple test) (Figure 4e). Results suggested that the wild-type genetic background but not the *w*^+^ locus reduced the number of nondirectional walks.

### w+ locus increased the maximal duration of one-way walking

The parameters of directional change and number of one-way or nondirectional walks were not very informative of how persistent a fly walked in one component. For instance, Canton-S and *w*^1118^ F10 flies might have different durations of one-way walking, although they had the same numbers of directional changes in the arenas. Through a careful control of genetic background, we examined the contribution of *w*^+^ locus to the maximal duration of one-way walking (MDOW).

It was common that during 300 s locomotion the MDOW were remarkably longer than those of nondirectional walking in each genotype of the tested flies (Figure 5). With the same genetic background, Canton-S had the MDOW (median 184.6 s, IQR 129.2 - 235.4 s, n = 31) longer than *w*^1118^ F10 flies (median 97.8 s, IQR 76.4 - 156.2 s, n = 43) (*P* = 0.0033, Mann-Whitney test). Clearly, Canton-S flies were able to walk in one direction, counter-clockwise or clockwise, for the maximal durations at a median of 184.6 s. Similarly, under the same genetic background, *w*^+^ F10 flies had the MDOW (median 31.0 s, IQR 18.2 - 36.4 s, n = 23) longer than *w*^1118^ flies (median 21.3 s, IQR 16.0 - 26.9 s, n = 32) (*P* = 0.0186, Mann-Whitney test). Thus, flies carrying *w*^+^ locus had increased MDOW compared with flies carrying *w*^1118^ locus. In contrast, *w*^+^/*w*^1118^ F1 female flies showed MDOW (median 35.2 s, IQR 27.3 - 43.5 s, n = 24) statistically the same as *w*^1118^/*w*^+^ F1 (median 32.9 s, IQR 17.0 - 49.3 s, n = 16) (*P* = 0.5520, Mann-Whitney test). Therefore, flies with identical genetic composition in the *w* allele and genetic background had statistically the same MDOW. These data indicated that *w*^+^ locus increased the maximal duration of one-way episodes in the circular arena.

**Figure 5:**
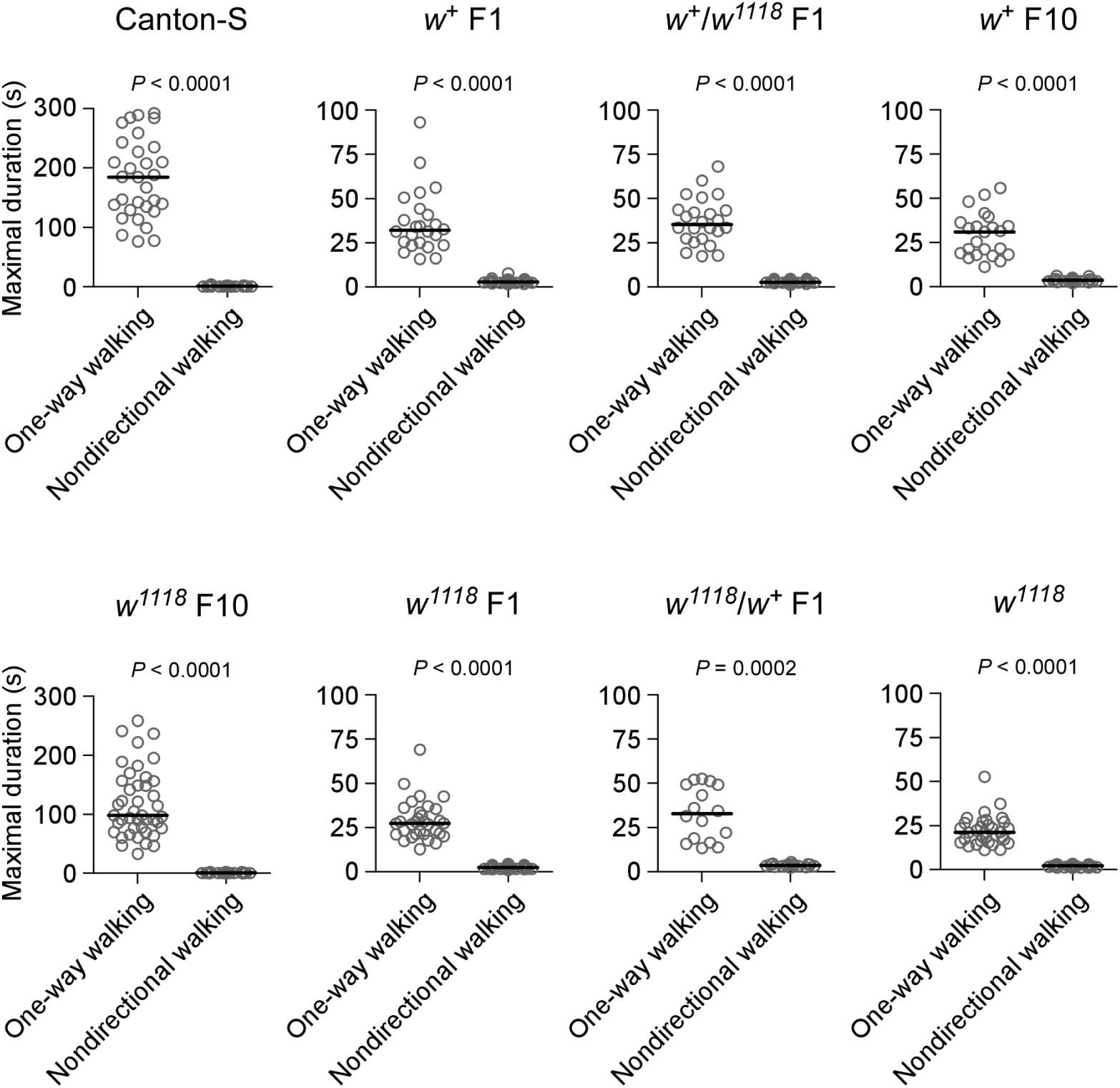
Maximal durations of walking components in tested flies. The maximal durations for one-way walking (including counter-clockwise and clockwise walking) and nondirectional walking during 300 s locomotion in tested flies. Fly genotypes are indicated. See the text for description of F1 and F10 flies. Data are presented as scattered dots and median (black line). *P* values from Mann-Whitney test.

### *w*^+^ duplicated to the Y chromosome increased the maximal duration of oneway walking

Direct evidence is required to support the hypothesis that the *w*^+^ gene promotes persistent one-way walking in circular arena. We examined flies with *w*^+^ duplicated to the Y chromosome. Two duplication lines (*w*^1118^/Dp(1;Y)*B*^*S*^*w*^+^*y*^+^ and *w*^1118^/Dp(1;Y)*w*^+^*y*^+^, both with isogenic background) have been previously established (Xiao and Robertson 2016; Xiao et al. 2017).

The MDOW in *w*^1118^/Dp(1;Y)*B*^*S*^*w*^+^*y*^+^ flies (median 32.3 s, IQR 24.2 - 41.0 s, n = 42) were longer than those in *w*^1118^ flies (*P <* 0.001, Kruskal-Wallis test with Dunn’s multiple comparison). The MDOW in *w*^1118^/Dp(1;Y)*w*^+^*y*^+^ flies (median 38.2 s, IQR 29.0 - 49.6 s, n = 47) were also longer than those in *w*^1118^ flies (*P <* 0.001, Kruskal-Wallis test with Dunn’s multiple comparison) (Figure 6a). Therefore, *w*^+^ duplicated to the Y chromosome increased the maximal duration of one-way episodes in circular arena.

**Figure 6:**
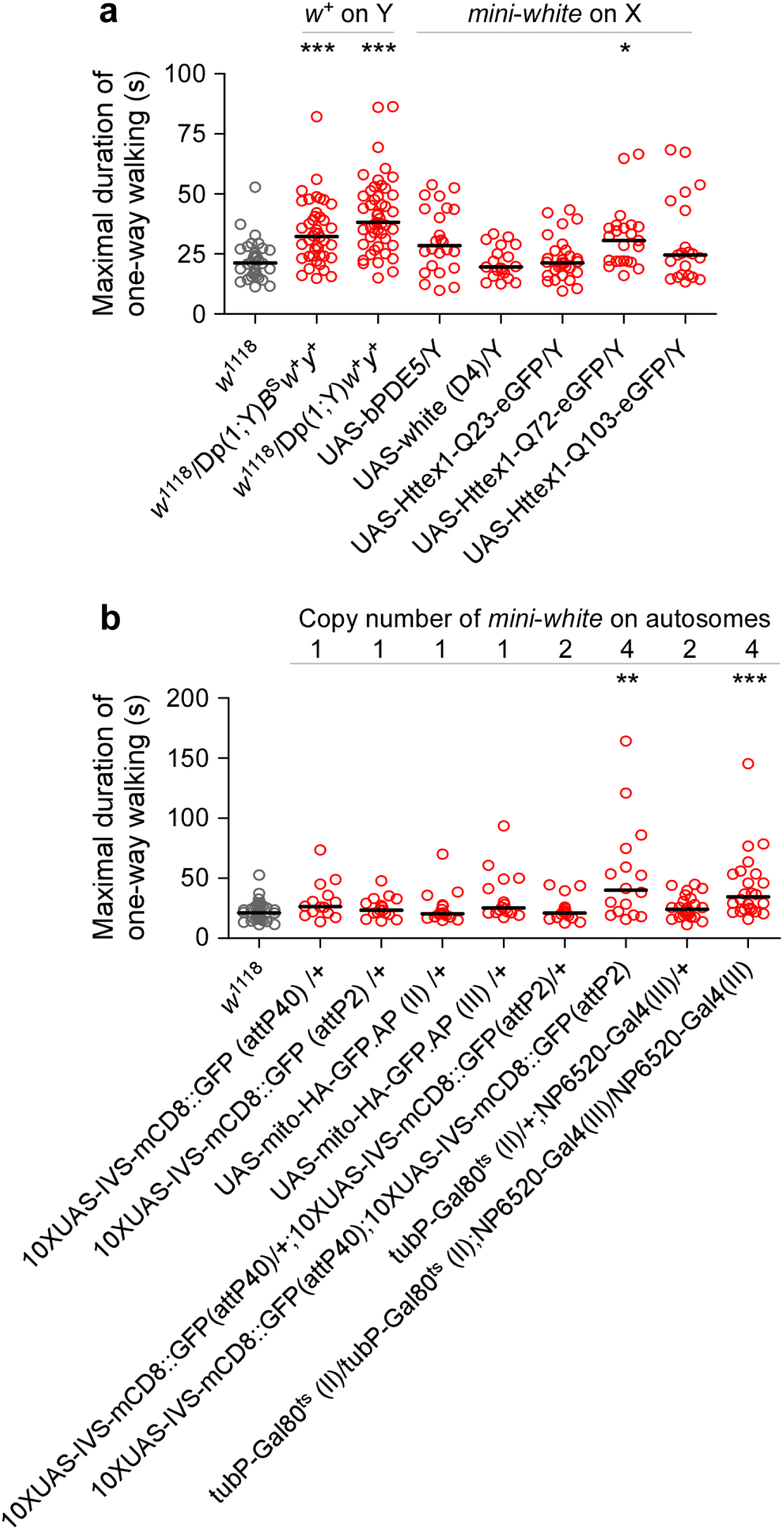
*w*^+^ increased the maximal duration of one-way walking. (**a**) Effect of *w*^+^ duplicated to the Y chromosome or *mini-white* inserted to the X chromosome on the maximal duration of one-way walking. (**b**) Effects of different copies of *mini-white* inserted to the autosomes on the maximal duration of one-way walking. Copy number of *mini-white* are listed. Data from *w*^+^ or *mini-white*-carrying flies are labeled with red. Data of *w*^1118^ flies (grey) are duplicated here from Figure 5 for comparison. *, *P <* 0.05; **, *P <* 0.01; ***, *P <* 0.001 from Kruskal-Wallis test with Dunn’s multiple comparison. Tested flies have been generated or backcrossed into isogenic background.

### Four copies of *mini-white* increased the maximal duration of one-way walking

The *mini-white* gene, a miniature form of *w*^+^, has been integrated into the genomes of many transgenic lines of *Drosophila* (e.g. UAS lines). These flies are ideal resources to test whether *w*^+^ promotes persistent one-way walking in circular arena.

We examined male flies carrying a *mini-white* on the X chromosome. All selected fly lines have been generated in *w*^1118^ isogenic background. Notably, the *mini-white* on the X chromosome is subject to a dosage compensation (Qian and Pirrotta 1995; Arkhipova et al. 1997). The MDOW were at a median of 28.5 s (IQR 19.5 - 43.8 s, n = 24) in UAS-bPDE5/Y, 19.6 s (IQR 15.8 - 28.8 s, n = 19) in UAS-*white* (D4)/Y, 21.3 s (IQR 17.1 - 26.5 s, n = 30) in UAS-Httex1-Q23-eGFP/Y, 30.6 s (IQR 22.0 - 36.4 s, n = 23) in UAS-Httex1-Q72-eGFP/Y and 24.6 s (IQR 15.7 - 45.2 s, n = 21) in UAS-Httex1-Q103-eGFP/Y. Male flies of UAS-Httex1-Q72-eGFP/Y showed increased MDOW compared with *w*^1118^ flies (*P <* 0.05, Kruskal-Wallis test with Dunn’s multiple comparison). There was no significant difference between each of the rest of tested flies and *w*^1118^ flies (*P >* 0.05, Kruskal-Wallis test with Dunn’s multiple comparison) (Figure 6a). Thus, a *mini-white* on the X chromosome was observed to be inconsistent to increase the maximal duration of one-way episodes.

Flies carrying one genomic copy of *mini-white* on the autosome were also examined. We chose UAS lines with *mini-white* integrated into site-specific recombination site (i.e. attP40 or attP2), and UAS lines with *mini-white* randomly inserted into the second or third chromosome. The selected UAS flies have been backcrossed into isogenic *w*^1118^ flies for ten generations. The MDOW were at a median of 23.5 s (IQR 20.1 - 32.1 s, n = 16) in 10*×*UAS-IVS-mCD8::GFP (attP40)/+ flies, 26.4 s (IQR 20.5 - 38.2 s, n = 14) in 10*×*UAS-IVS-mCD8::GFP (attP2)/+ flies, 20.6 s (IQR 17.7 - 27.6 s, n = 16) in UAS-mito-HA-GFP.AP (II)/+ flies and 25.3 s (IQR 21.3 - 47.4 s, n = 16) in UAS-mito-HA-GFP.AP (III)/+ flies. There was no statistical difference between each of the tested UAS flies and *w*^1118^ flies (Kruskal-Wallis test with Dunn’s multiple comparison). Thus, flies with one genomic copy of *mini-white* on the autosome showed unaffected MDOW (Figure 6b).

Flies containing two or four genomic copies of *mini-white* on the autosomes were tested. Flies carrying two genomic copies of *mini-white* (10*×*UAS-IVS-mCD8::GFP (attP40)/+;10*×*UAS-IVS-mCD8::GFP (attP2)/+) had the MDOW at a median of 21.1 s (IQR 17.1 - 25.7 s, n = 16), a level comparable with that in *w*^1118^ flies (*P >* 0.05, Kruskal-Wallis test with Dunn’s multiple comparison). Flies carrying four genomic copies of *mini-white* (10*×*UAS-IVS-mCD8::GFP (attP40);10*×*UASIVS-mCD8::GFP (attP2) homozygotes) showed the MDOW at a median of 40.0 s (IQR 20.1 - 71.1 s, n = 16), a level higher than that in *w*^1118^ flies (*P <* 0.01, Kruskal-Wallis test with Dunn’s multiple comparison). Consistently, another line of flies carrying two genomic copies of *mini-white* (tubP-Gal80^*ts*^/+;NP6520-Gal4/+) had the MDOW (median 24.2 s, IQR 18.0 - 32.6 s, n = 24) comparable with *w*^1118^ flies (*P >* 0.05, Kruskal-Wallis test with Dunn’s multiple comparison), while flies with four genomic copies of *mini-white* (tubP-Gal80^*t*^*s*;NP6520-Gal4 homozygotes) had increased MDOW (median 34.5 s, IQR 24.2 - 53.6 s, n = 24) compared with *w*^1118^ flies (*P <* 0.001, Kruskal-Wallis test with Dunn’s multiple comparison) (Figure 6b).

Together, these data indicated that four genomic copies of *mini-white*, if integrated into autosomes as two homozygous alleles, increased the maximal duration of one-way walking in circular arena, whereas one copy of *mini-white* on the X chromosome, or one or two copies on the autosomes had no or inconsistent effect.

### Pan-neuronal overexpression of *w*^+^ increased the maximal duration of oneway walking

Often, the Gal4*/*UAS expression system introduces two genomic copies of *mini-white*, which if integrated into autosomes, were observed to be insufficient to increased the MDOW. This validated the application of Gal4*/*UAS system to explore the effect of tissue-specific expression of *w*^+^ on persistent one-way walking. Presumably, overexpression of the White protein in targeted tissues, for example, the central nervous system, could have an effect on the MDOW. We examined flies with targeted expression of *w*^+^ using pan-neuronal driver elav-Gal4.

Flies expressing White in the neurons (elav-Gal4/+;UAS-*white* (H8)/+) had the MDOW (median 40.5 s, IQR 23.2 - 53.3 s, n = 12) longer than elav-Gal4/+ (median 19.8 s, IQR 17.9 - 22.6 s, n = 16) (*P <* 0.05, Kruskal-Wallis test with Dunn’s multiple comparison) or UAS-*white* (H8)/+ (median 25.0 s, IQR 17.7 - 27.4 s, n = 16) (*P <* 0.05, Kruskal-Wallis test with Dunn’s multiple comparison). Flies expressing an irrelevant protein, the green fluorescent protein (GFP), in the neurons (elav-Gal4/+; 10*×*UAS-IVS-GFP-WPRE (attP2)/+) had the MDOW (median 26.8 s, IQR 18.6 - 41.3 s, n = 8) the same as controls 10*×*UAS-IVS-GFP-WPRE (attP2)/+ (median 24.3 s, IQR 21.3 - 28.8 s, n = 8) (*P >* 0.05, Kruskal-Wallis test with Dunn’s multiple comparison) (Figure 7a). Thus, pan-neuronal overexpression of the White protein but not GFP increased the maximal duration of one-way walking in circular arena.

**Figure 7:**
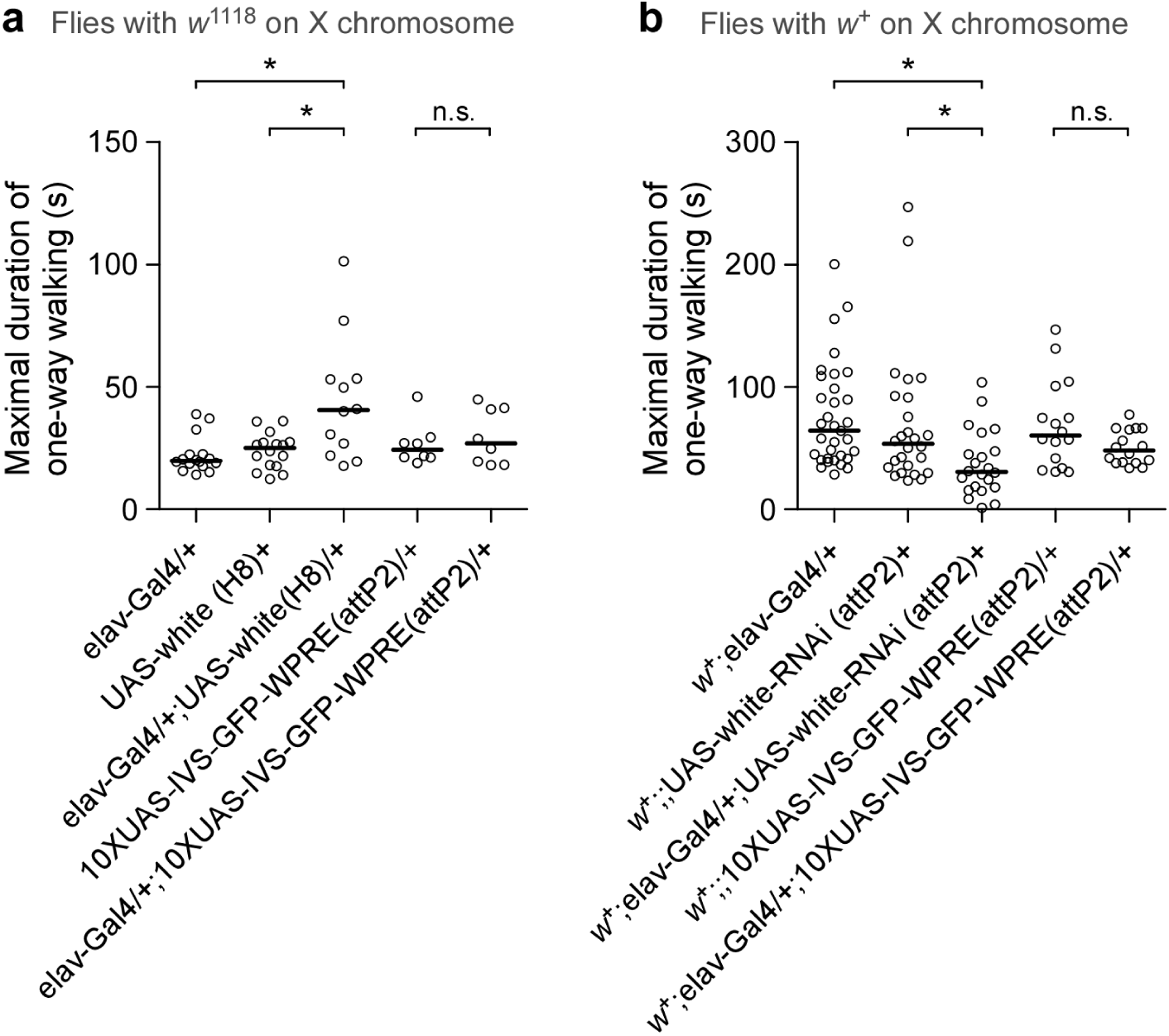
Pan-neuronal overexpression or downregulation of *w*^+^ affects the maximal duration of one-way walking. (**a**) Pan-neuronal overexpression of *w*^+^ increases the maximal duration of one-way walking. There is no effect of Gal4/UAS-driven overexpression of GFP. Tested flies carry a *w*^1118^ allele on the X chromosome. (**b**) Pan-neuronal knockdown of *w*^+^ through RNAi decreases the maximal duration of one-way walking. There is no effect of targeted expression of GFP by the same system. Tested flies carry a *w*^+^ on the X chromosome. *, *P <* 0.05 from Kruskal-Wallis test with Dunn’s multiple comparison. n.s., non-significance.

### RNAi knockdown of *w*^+^ decreased the maximal duration of one-way walking

Downregulation of *w*^+^ through RNA interference (RNAi) provided another approach to examine the effect of *w*^+^ on persistent one-way walking. After a recombination of wild-type X chromosome (including target gene *w*^+^) into elav-Gal4 and UAS-*white*-RNAi flies, we examined the effect of RNAi knockdown of *w*^+^ on the MDOW.

The MDOW in *white-RNAi* flies (*w*^+^; elav-Gal4/+; UAS-*white*-RNAi (attP2)/+, median 30.8 s, IQR 17.6 - 51.3 s, n = 22) were decreased compared with controls *w*^+^; elav-Gal4/+ (median 64.2 s, IQR 41.6 - 103.7 s, n = 33) (*P <* 0.05, Kruskal-Wallis test with Dunn’s multiple comparison) or controls *w*^+^; UAS-*white*-RNAi (attP2)/+ (median 53.5 s, IQR 33.1 - 91.7 s, n = 26) (*P <* 0.05, Kruskal-Wallis test with Dunn’s multiple comparison). The MDOW in flies expressing a GFP (*w*^+^; elav-Gal4/+; 10*×*UAS-IVS-GFP-WPRE (attP2)/+, median 48.2 s, IQR 38.2 - 66.0 s, n = 16) was statistically the same as controls *w*^+^; 10*×*UAS-IVS-GFP-WPRE (attP2)/+ (median 60.4 s, IQR 35.7 - 94.2 s, n = 16) (*P >* 0.05, Kruskal-Wallis test with Dunn’s multiple comparison) (Figure 7b).

Therefore, RNAi knockdown of *w*^+^ decreased the maximal duration of one-way walking. Without targeted RNAi to *w*^+^, targeted expression of GFP using the same system had no effect. Data supported that *w*^+^ promoted persistent one-way walking in circular arena.

## Discussion

*Drosophila* persistent one-way walking in a circular arena is a phenomenon that has not been previously reported. Using the techniques of fly tracking, behavioral computation and genetic manipulation, we show that wild-type Canton-S male flies are able to walk in one direction, counterclockwise or clockwise, for a median value around 185 s in the circular arenas. Whereas the genetic background of Canton-S reduces directional change and numbers of one-way and nondirectional walks, *w*^+^ promotes persistent one-way walking by increasing the maximal duration of counter-clockwise or clockwise walking.

### Features of persistent one-way walking

The extraction of walking elements into four quantifiable components, including counter-clockwise walking, clockwise walking, nondirectional walking and pausing, provides a foundation for the characterization of complicated walking behavior in *Drosophila*. Canton-S and *w*^1118^ male flies increase locomotion in the circular arenas as a response to spatial restriction, and maintain active walking for at least an hour (Xiao and Robertson 2015). Here we demonstrate further details of walking behavior in addition to the increased locomotion. We summarize our observations that are common to Canton-S and *w*^1118^ flies from three aspects.

First, counter-clockwise walking and clockwise walking are the two primary components comprising the largest time proportion. There is no preference for counter-clockwise or clockwise direction in the arena. Second, intermittent pausing is predominantly associated with directional persistence but not directional change. Thus, pausing is indicative of a state that flies rest and retain a memory of walking direction, rather than a state in which flies are unsure of their directions. Cocaine-treated Canton-S flies show a general tendency to rotate in one direction both before and after an immobility (Gomez-Marin et al. 2016), a finding similar to our observation that Canton-S flies have a strong tendency to walk in the same direction both before and after a pause in circular arenas. Third, during counter-clockwise or clockwise walking flies move forward. We did not observe flies walking backward persistently in the arenas. Therefore, persistent one-way walking is different from the phenotype of walking backward consistently (Bidaye et al. 2014).

### Effects of wild-type genetic background on persistent one-way walking

Canton-S and *w*^1118^ flies have two major genetic differences: the *w* allele and its genetic background. The effect of cytoplasmic background on persistent one-way walking is negligible, because female progenies from reciprocal crosses between Canton-S and *w*^1118^ flies have the same directional change of walking. The observations that low number of directional changes, and low numbers of one-way walks and nondirectional walks are transferable, from wild-type to *w*^1118^-carrying flies with wild-type genetic background, suggest that wild-type genetic background greatly reduces directional change and the numbers of one-way and nondirectional walks in circular arena. Additionally, a 1:1 mixture of genomic content (including *w* allele and genetic background) between Canton-S to *w*^1118^ flies removes the difference of directional change of walking. Even F1 male flies with a half-half mixture of autosomes, despite the presence of wild-type or isogenic X chromosome, have the same numbers of directional changes. These findings highlight that wild-type genetic background is strongly associated with persistent one-way walking. Thus, to examine the contribution of *w*^+^ to this phenotype, the genetic background should be carefully controlled.

### *w*^+^ promotes persistent one-way walking

The *w*^+^ locus has no effect on the directional change, the number of one-way walks or the number of nondirectional walks. Thus, the *w*^+^ locus and wild-type genetic background might have respectively specific effects in promoting persistent one-way walking. With the same genetic background, flies carrying *w*^+^ locus have increased MDOW compared with flies carrying *w*^1118^ locus. This finding suggests that the *w*^+^ gene promotes persistent one-way walking by increasing the maximal duration of one-way episodes.

Further observations have supported this suggestion. The first is that *w*^+^ duplicated to the Y chromosome increases the MDOW in flies with isogenic background. Second, four genomic copies (or two homozygous alleles) of *mini-white* on the autosomes increase the MDOW, whereas a single copy on the X chromosome, or one or two copies on the autosomes have no or inconsistent effect. Likely, four genomic copies of *mini-white* results in an increased expression of the White protein, and thus an observable effect. This copy-number-dependent phenomenon has been observed elsewhere. It has been reported that *mini-white* is responsible for another complex behavior, the male-female copulation success, in a manner that is copy-number-dependent (Xiao et al. 2017).

It is justified to apply the *mini-white*-carrying Gal4*/*UAS expression system if less than four genomic copies of *mini-white* are introduced into autosomes, to explore the effect of tissue-specific overexpression of the White protein on persistent one-way walking. Pan-neuronal expression of White increases the MDOW. There is, however, no effect through the expression of an irrelevant protein GFP instead of White using the same Gal4*/*UAS system. These findings support that *w*^+^ promotes persistent one-way walking, and that two genomic copies of *mini-white* introduced by the Gal4*/*UAS system have no effect. The effect of *w*^+^ is further supported by the observations that RNAi knockdown of *w*^+^ reduces the maximal duration of one-way walking, and that there is no effect by targeted expression of GFP without a knockdown of *w*^+^ using the same expression system. Altogether, these observations confirm that *w*^+^ promoted persistent one-way walking by increasing the maximal duration of one-way episodes.

It is worth noting that the suspected influence of visual ability is negligible, because Canton-S flies under dim red illumination, in which flies are virtually blind due to the poor visual sensitivity (McEwen 1918), have regular walking trajectories similar to flies with white light illumination.

It is intriguing that in general, a Canton-S fly is able to walk in one direction, counterclockwise or clockwise, for around 185 seconds in a circular arena, while a *w*^1118^ fly only 21 seconds. We have previously reported that Canton-S flies have spontaneously rhythmic motor activities with a periodicity around 19 s, but this periodicity is largely diminished in *w*^1118^ flies (Qiu et al. 2016).Although it has been shown that the *w*^+^ gene increases the consistency of this periodicity, whether such periodic motor activities promote persistent one-way walking in Canton-S flies remains elusive.

For over a hundred years, it has been believed that *w*^+^ is responsible for eye color in *Drosophila*. This has led to an application of *mini-white* as a marker indicating the successful transformation of a transgene. This application heavily relies on the supposition that *w*^+^ has an exclusive role in eye pigmentation. However, *mini-white* has additional functions in causing male-male courtship chaining (Zhang and Odenwald 1995; Hing and Carlson 1996), and conferring male-female copulation success (Xiao et al. 2017). Furthermore, *w*^+^ promotes fast and consistent locomotor recovery from anoxia (Xiao and Robertson 2016, 2017). Wild-type flies have enhanced memory of thermal stimulation (Sitaraman et al. 2008), and increased vesicular content of biogenic amines (Borycz et al. 2008) compared with *w* mutant flies. The extra-retinal function that the White protein transports cyclic guanosine monophosphate (cGMP) has been documented (Evans et al. 2008; Xiao and Robertson 2017). These findings suggest that *w*^+^ has pleiotropic housekeeping functions rather than a function exclusively responsible for eye color. These housekeeping functions could be essential to promote persistent one-way walking in a circular arena.

In summary, the findings that Canton-S flies are able to walk in one direction in circular arenas for around 185 seconds, and that *w*^+^ increases the maximal duration of one-way walking firmly support a pleiotropic function of *w*^+^ in promoting persistent one-way walking.

## Statement of human and animal rights

This article does not contain any studies with human participants performed by any of the authors.

## Supplementary material

Video S1: Walking activities of Canton-S flies in the circular arenas

Video S2: Walking activities of *w*^1118^ flies in the circular arenas

---

**Figure S1:**
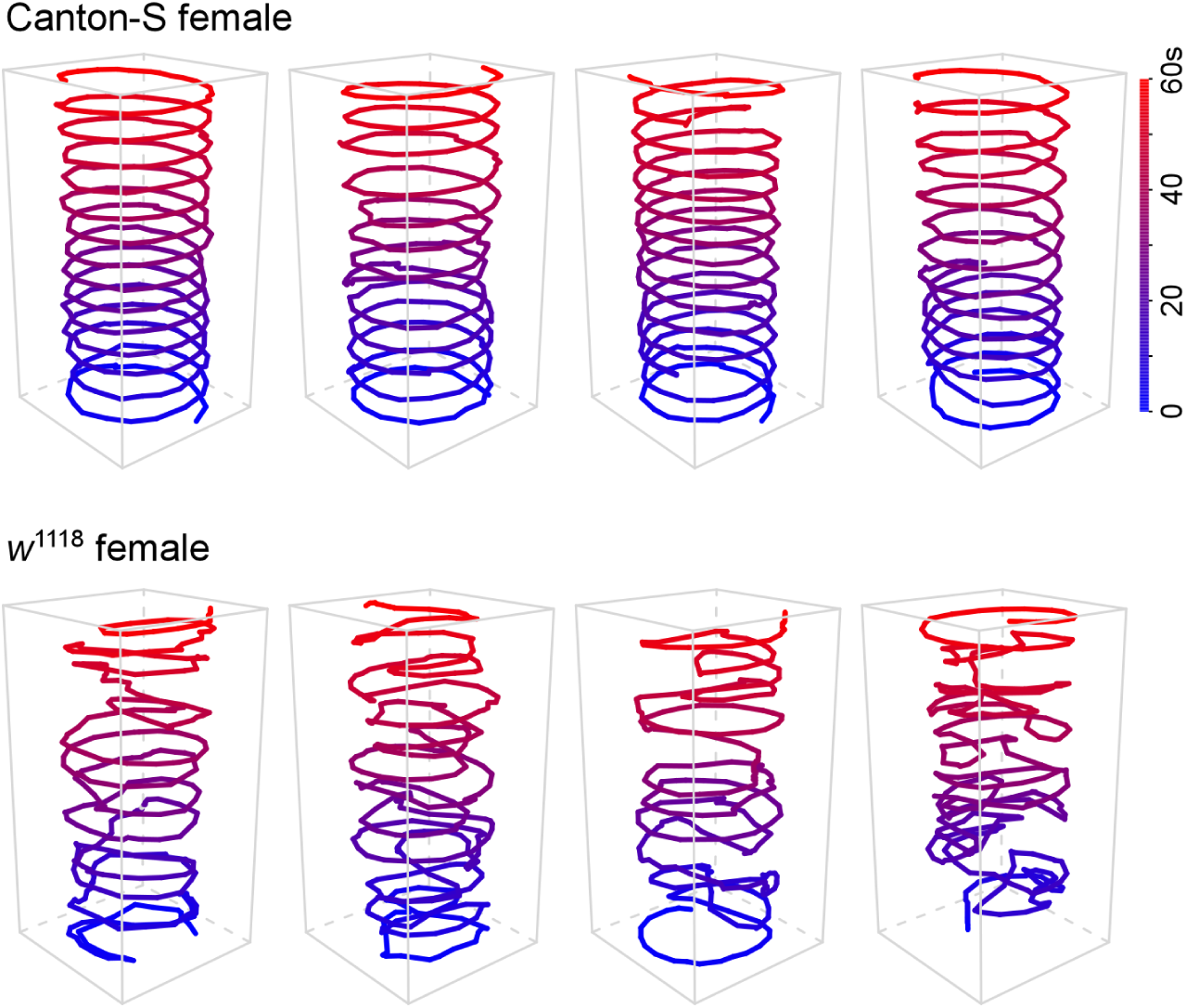
Walking trajectories in Canton-S and *w*^1118^ female flies. Walking trajectories in circular arenas were observed to be visually regular in Canton-S (upper panel), but irregular in *w*^1118^ (lower panel) female flies. For clarity, activities from only 60 s are shown. Color key indicates the time.

**Figure S2:**
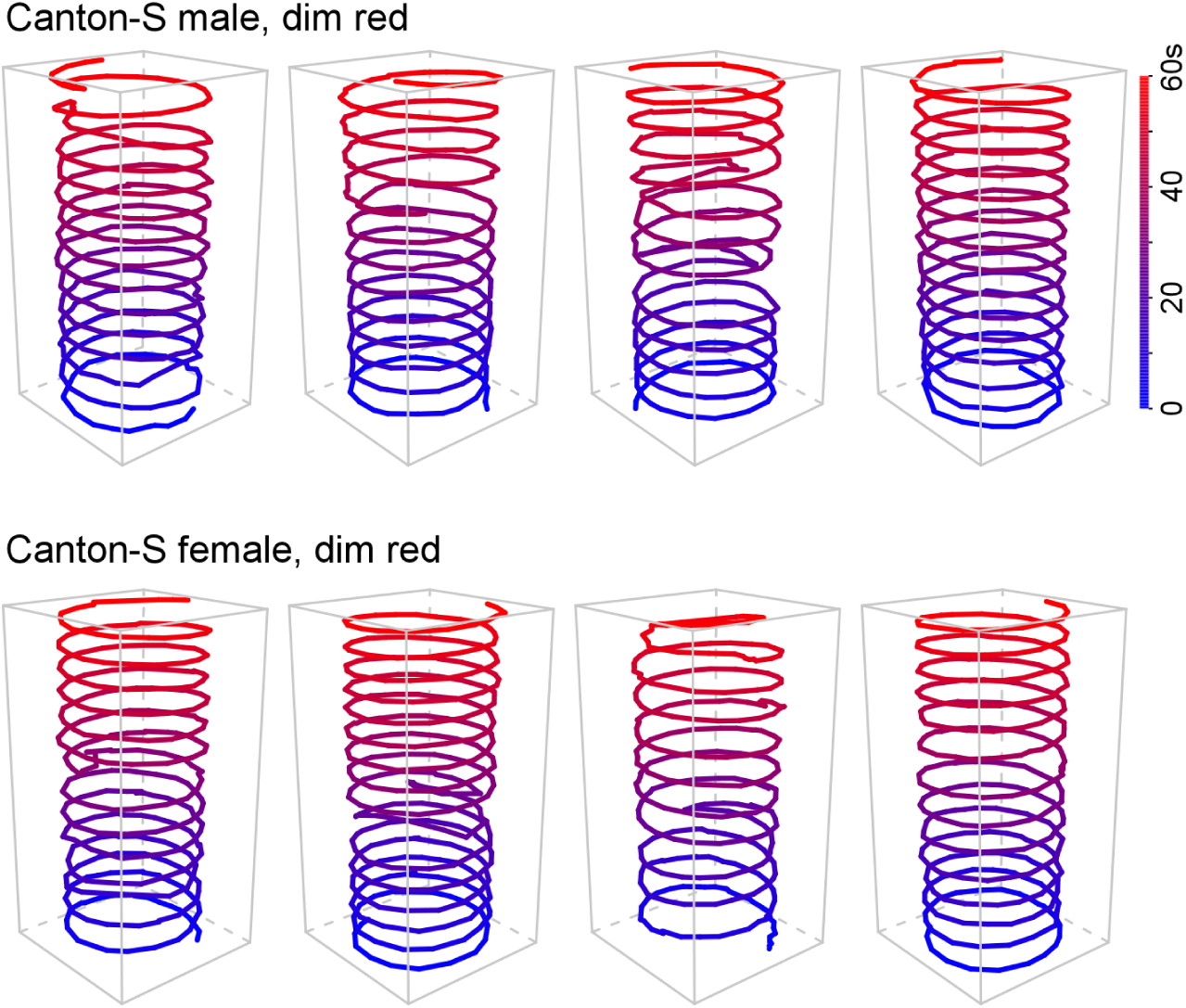
Walking trajectories of Canton-S with dim red illumination. Persistent one-way walking in Canton-S flies was unaffected by dim red illumination. For clarity, activities from only 60 s are shown. Color key indicates the time.

**Figure S3:**
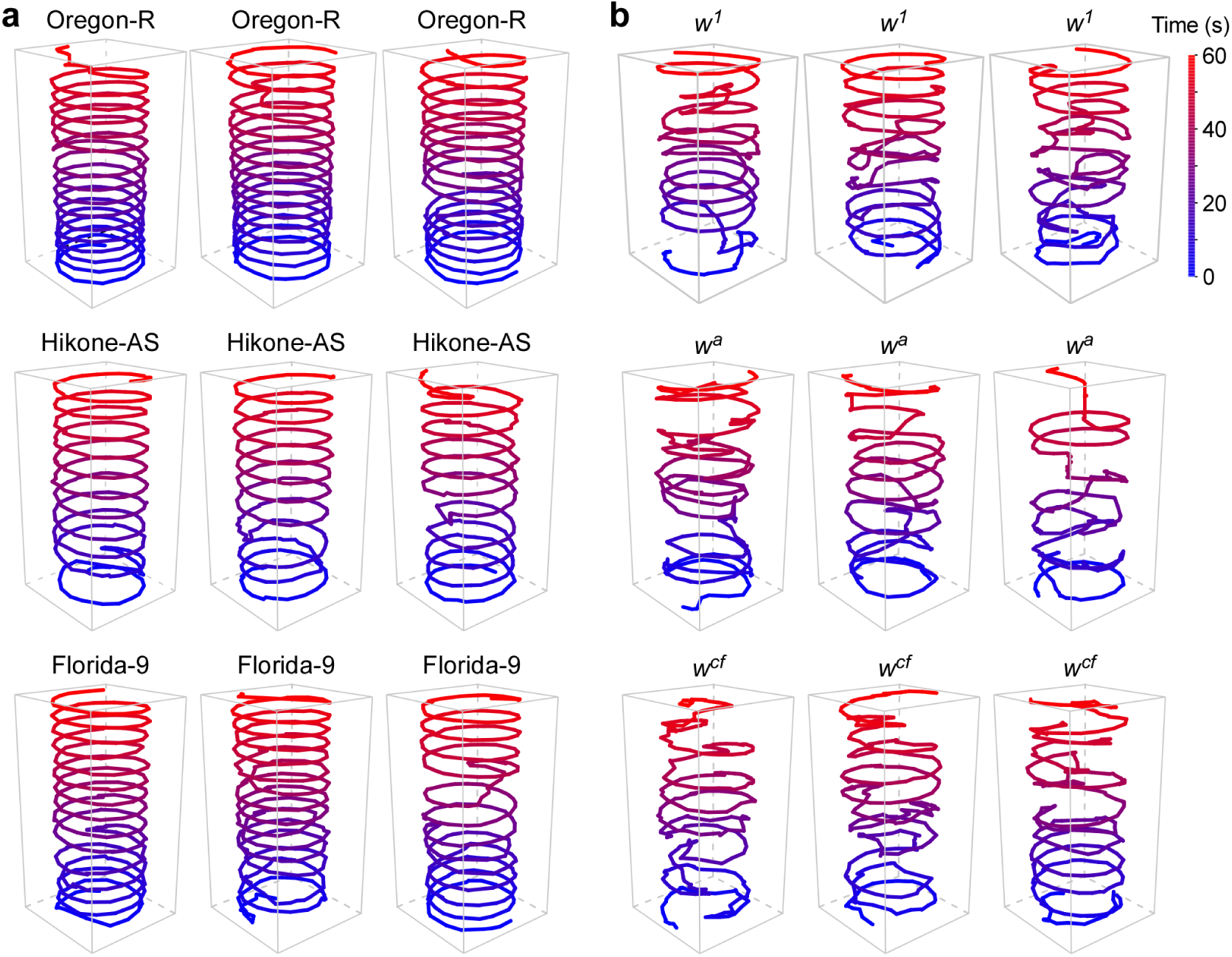
Walking trajectories of several wild-types and *w* mutants. (**a**) Walking trajectories of Oregon-R, Hikone-AS and Florida-9 in the circular arenas. (**b**) Walking trajectories of *w*^1^, *w*^*a*^and *w*^*cf*^ in the circular arenas. For clarity, activities from only 60 s are shown. Color key indicates the time.

